# Linking meta-omics to the kinetics of denitrification intermediates reveals pH-dependent causes of N_2_O emissions and nitrite accumulation in soil

**DOI:** 10.1101/2020.11.26.399899

**Authors:** Åsa Frostegård, Silas HW Vick, Natalie YN Lim, Lars R Bakken, James P Shapleigh

## Abstract

Denitrifier community phenotypes often result in transient accumulation of denitrification (NO_3_^−^→NO_2_^−^→NO→N_2_O→N_2_) intermediates. Consequently, anoxic spells drive NO^-^, N_2_O^-^ and possibly HONO^-^emissions to the atmosphere, affecting both climate and tropospheric chemistry. Soil pH is a key controller of intermediate levels, and while there is a clear negative correlation between pH and emission of N_2_O, NO_2_^−^ concentrations instead increase with pH. These divergent trends are probably a combination of direct effects of pH on the expression/activity of denitrification enzymes, and an indirect effect via altered community composition. This was studied by analyzing metagenomics/transcriptomics and phenomics of two soil denitrifier communities, one of pH 3.8 (Soil3.8) and the other 6.8 (Soil6.8). Soil3.8 had severely delayed N_2_O reduction despite early transcription of *nosZ*, encoding N_2_O reductase, by diverse denitrifiers, and of several *nosZ* accessory genes. This lends support to a post-transcriptional, pH-dependent mechanism acting on the NosZ apo-protein or on enzymes involved in its maturation. Metagenome/metatranscriptome reads of *nosZ* were almost exclusively clade I in Soil3.8 while clade II dominated in Soil6.8. Reads of genes and transcripts for NO_2_^−^-reductase were dominated by *nirK* over *nirS* in both soils, while qPCR-based determinations showed the opposite, demonstrating that standard primer pairs only capture a fraction of the *nirK* community. The -omics results suggested that low NO_2_^−^ concentrations in acidic soils, often ascribed to abiotic degradation, are primarily due to enzymatic activity. The NO reductase gene *qnor* was strongly expressed in Soil3.8, suggesting an important role in controlling NO. Production of HONO, for which some studies claim higher, others lower, emissions from NO_2_^−^ accumulating soil, was estimated to be ten times higher from Soil3.8 than from Soil6.8. The study extends our understanding of denitrification-driven gas emissions and the diversity of bacteria involved and demonstrates that gene and transcript quantifications cannot always reliably predict community phenotypes.

## Introduction

During the past century human activities have accelerated the input of reactive N to the biosphere [1, 2]. This has escalated the emissions of N_2_O, a major greenhouse gas and contributor to ozone depletion [3, 4, 5], as well as nitric oxide (NO) and nitrous acid (HONO) which both influence chemical reactions in the troposphere, leading to formation of undesired ozone [6]. Agriculture, and especially the excessive use of synthetic N fertilizers in large parts of the world, is a major source of N_2_O release from terrestrial systems, primarily via denitrification and nitrification [1, 7, 8]. While CO_2_ emissions are predicted to decline substantially during the present century, anthropogenic N_2_O emissions, which have increased at an average rate of 0.6 ± 0.2 Tg N yr^−1^ per decade during the past 40 years, are expected to continue to increase [5] or remain nearly constant [9], unless new methods to reduce the N_2_O/N_2_ product ratio of our agroecosystems are developed [10, 11]. Some mitigation options have been identified through improved understanding of the organisms producing and reducing N_2_O and the environmental factors that control these emissions [12], and more are expected to emerge as we intensify our research on denitrification and denitrifying organisms in soil.

Denitrification is the stepwise reduction of nitrate (NO_3_^−^) to N_2_ via nitrite (NO_2_^−^), NO and N_2_O, and is performed by a diverse range of facultatively anaerobic organisms when O_2_ becomes scarce. Complete reduction of NO_3_^−^ to N_2_ is a four-step process with each step catalysed by a different reductase [13, 14]: the NO_3_^−^ reductases NarG (membrane-bound) and/or NapA (periplasmic) encoded by the genes *narG* and *napA*; the NO_2_^−^ reductases NirK (copper-containing) or NirS (containing cytochrome *cd*1) encoded by *nirK* and *nirS*; the NO reductases cNor (cytochrome *c* dependent) or qNor (quinol-dependent) encoded by *cnor* and *qnor*; and the N_2_O reductase NosZ, of which two clades have been identified, encoded by *nosZ* clade I and II [15]. While many organisms have both Nar and Nap, most denitrifiers carry only one type of Nir, Nor and NosZ. The denitrification pathway is modular [16] and the absence of one or more of the reduction steps is common [17, 18]. Not only does the presence/absence of the denitrification genes affect the phenotypes of denitrifiers and the amounts of the different denitrification products that are released but gene regulation [14], in interplay with environmental factors, is a key factor as well [18, 19, 20]. Similar controls of N_2_O/N_2_ product ratios have been observed in soil communities [21]. If the production and reduction steps are not fully balanced net production of a denitrification intermediate will occur. Studies of denitrification kinetics during anoxic incubation of various denitrifying organisms always show temporal accumulation of intermediate products [18, 22], but the extent differs even between closely related strains [23]. Also, since the timing of denitrification gene transcription varies between organisms, denitrifier communities will have a succession of actively transcribing populations during an anoxic spell which will also influence levels of intermediate products [24].

Of the environmental key factors known to control denitrification perhaps the best characterized is soil pH [25, 26], which profoundly affects the accumulation of denitrification intermediates. The effects of pH on the regulation and enzymology of the reduction steps differ, however, and the mechanisms by which pH impacts these processes are not well understood. While the accumulation of NO_2_^−^ increases with soil pH [27, 28], there is a clear negative correlation between pH and N_2_O emissions [29–33]. The simple assumption that soils with high N_2_O emissions have fewer N_2_O^-^reducing organisms is corroborated by some studies but not by others [32, 34–36], which lends little support for a direct, causal relationship between *nosZ* numbers and N_2_O emissions. Studies of denitrifying bacteria in pure culture, extracted soil bacterial communities and in intact soils [37–39] all showed that *nosZ* transcription did take place under acidic conditions (pH 5.7-6.1) during periods of active denitrification but with negligible or strongly delayed N_2_O reduction. This suggests a post-transcriptional phenomenon, possibly interference with the maturation of the NosZ apo-protein, or with the transfer of electrons to the reductase. The quantification of *nosZ* transcripts in those soil communities was, however, based on PCR primers targeting only bacteria carrying *nosZ* clade I. Moreover, the transcripts were not sequenced, which leaves it open to speculation whether different fractions of the denitrifying bacteria transcribe *nosZ* at low versus high pH. This, and the fact that the post-transcriptional effect of low pH has only been tested in a few model strains, implies that it remains essentially unknown how widespread this phenomenon is among denitrifiers in soils.

The general occurrence of low NO_2_^−^ concentrations in acidic soils has been attributed mainly to chemical decomposition at low pH [40]. This was challenged recently by [28], who suggested that high rates of microbial reduction of NO_2_^−^ played a key role in keeping NO_2_^−^ concentrations low during denitrification in acidic soil. This would also imply high rates of NO reduction since the concentration of this gas was kept low. In soil with near neutral pH, on the other hand, the rate of NO_3_^−^ reduction exceeded the rate of NO_2_^−^ reduction, leading to NO_2_^−^ accumulation. The *nir* and *nor* gene abundance, their transcriptional activity and the organisms involved were however not investigated.

Here we aimed at resolving some of the questions raised from the above-mentioned studies about how soil pH affects the composition and activity of the denitrifier community and the accumulation/release of denitrification intermediates. We took an integrated “multi-omics” approach [41] to avoid primer biases [42] and thus to include as large a portion as possible of the microbial community, and analyzed the metagenomes (MGs) and the metatranscriptomes (MTs) of two soils with significantly different pH, 3.8 and 6.8. Samples for MT analysis were taken at time intervals through anoxic incubation of soil microcosms, during which the dynamics of the denitrifier community “phenome” was followed by frequent determinations of denitrification intermediate and end products. The results from the phenomics analysis, presented in more detail in [28], showed complete denitrification in Soil6.8 from the start of the incubation and accumulation of NO_2_^−^. Soil3.8, on the other hand, showed no NO_2_^−^ accumulation but severely delayed N_2_O reduction (Fig. 1). The problems with obtaining mRNA from low pH soil (pH 4) encountered earlier [38] were overcome using an optimized protocol [43], which provided nucleic acid quality suitable for both shotgun MG- and MT sequencing. The relative abundances of detected denitrification genes and transcripts were compared to results from PCR-quantification, which confirmed the severe biases related to amplification-based analyses.

**Figure 1.**
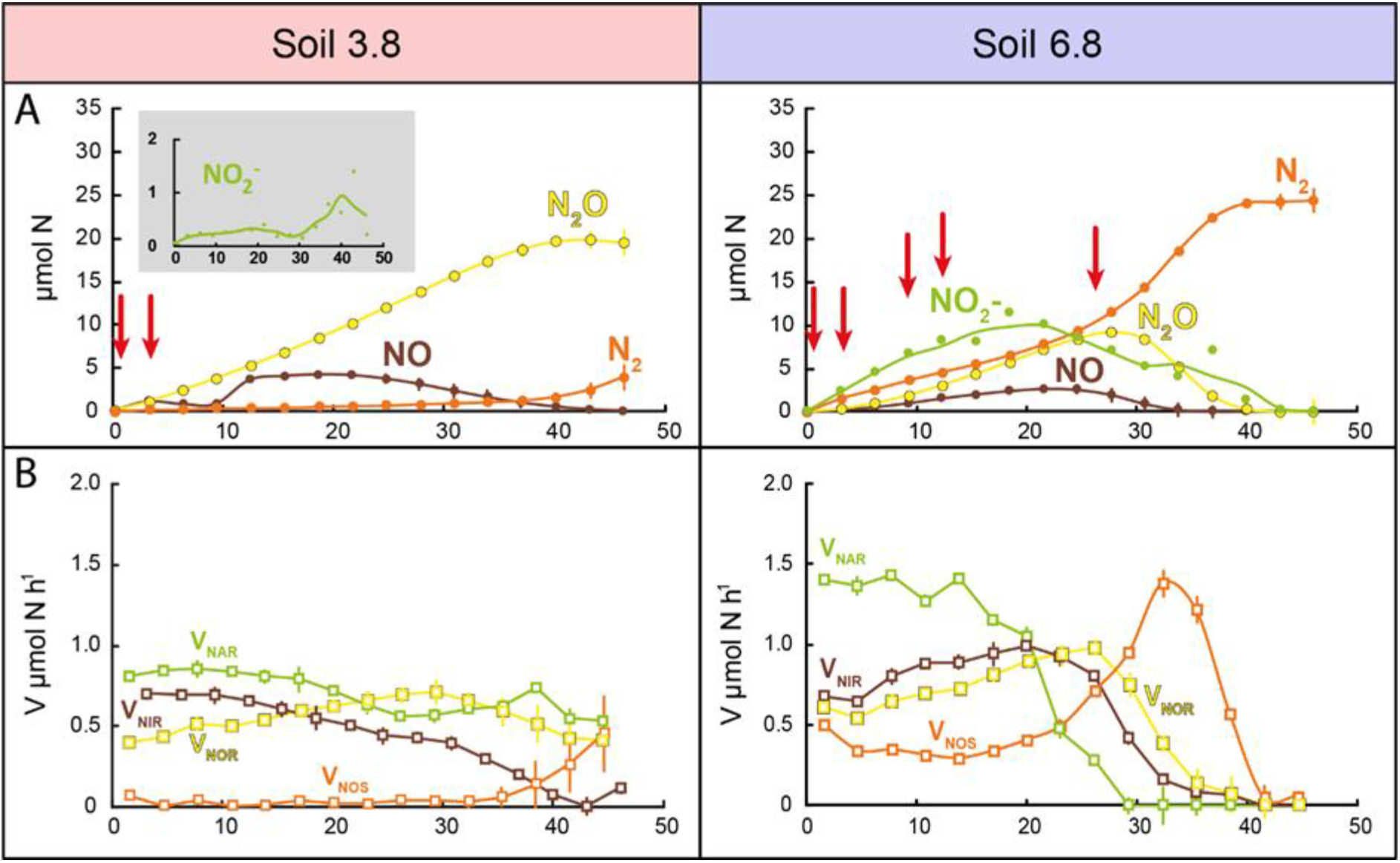
Kinetics of NO_2_^−^ and N-gases, calculated enzyme rates, and kinetics of NO_2_^−^ and HNO_2_ during anoxic incubation of the soil with pH_CaCl2_ = 3.8 and 6.8, amended with NO_3_^−^. **Panel A** shows the measured amounts of NO_2_^−^, NO, N_2_O and N_2_. Sampling for metatranscriptome analyses are indicated by red arrows (0.5 and 3 h for Soil3.8; 0.5, 3, 9, 12 and 27 h for Soil6.8). **Panel B** shows the reduction rates for the different denitrification steps *V*_*NAR*_ (NO_3_^−^→ NO_2_^−^), *V*_*NIR*_ (NO_2_^−^→NO), *V*_*NOR*_ (NO→N_2_O) and V_*NOS*_ (N_2_O→N_2_), all given as μ mol N vial^−1^ h^−1^. The values were based on the measured kinetics of NO_2_^−^, NO, N_2_O and N_2_ and corrected for abiotic decomposition of NO_2_^−^ as published previously by Lim *et al.* (2018). Abiotic NO_2_^−^ decomposition was significant only in Soil3.8. The figure is adapted from graphs shown in Lim et al., 2018, based on the same data set.

The -omics analyses allowed us to address several issues. One was to determine if the abundance of *nar/nap*, *nir* and *nor* genes and transcripts could explain the strong control of NO_2_^−^ and NO observed in the acidic soil and which organisms were involved. In addition, the detailed kinetics data made it possible to estimate if NO_2_^−^-accumulation in neutral soil would lead to more or less HONO emission than from non-NO_2_^−^ accumulating, acidic soil. Secondly, we investigated if the apparent lack of DNRA activity (dissimilatory nitrite reduction to ammonium) in these soils was due to low abundance of DNRA-related genes and transcripts. Thirdly, we clarified the complex ecophysiology of *nosZ* carrying bacteria to better understand their hampered N_2_O reduction under acidic conditions (this study and [38, 39]). To do so, we investigated to what extent the two *nosZ* clades were found in the MGs and MTs of two soils of differing pH; if *nosZ* gene transcripts originated from a few populations or represented diverse denitrifying bacteria; and if genes other than *nosZ* in the *nos* operon were transcribed in acidic soil. For the latter, we included *nosR*, which encodes NosR suggested to be involved in electron delivery to NosZ in organisms with *nosZ* cladeI; *nosL*, which encodes a chaperone delivering Cu to the NosZ apo-protein (both *nosZ* clades); and the ORF *nosDFY* (both *nosZ* clades) encoding NosD, suggested to be involved in NosZ maturation, and the ABC-transporter NosFY [44–46].

## Materials and methods

### Soils

Two peat soils (40-45 % organic C, 2 % organic N) [38] with pH 3.80 (Soil3.8) and 6.80 (Soil6.8) were sampled from a long-term field experimental site in western Norway (61°17’42“, 5°03’03”). Soil3.8 is the original un-limed soil, and Soil6.8 was limed in 1978 with shell sand (800 m^3^ ha^−1^; [47]). Freshly sampled soils were transported to the laboratory, sieved (4.5 mm) upon arrival, then stored in sealed plastic bags at 4 °C. All pH values were measured in 0.01 M CaCl_2_ (soil:CaCl_2_ 1:5) immediately prior to further analyses.

### Soil treatment

The soils were revitalised from cold storage by addition of 5 mg dried, powdered clover g^−1^ soil wet weight (ww), then incubated at 15 °C for 72 h [38]. Soil aliquots corresponding to 1.5 g soil organic C (5-8 g of soil ww depending on lime content) were placed in air-tight glass vials and sealed with butyl-rubber septa and aluminium crimps. Nitrate (as KNO_3_) was dissolved in autoclaved MilliQ water which was added to reach 80% of the soil’s water holding capacity (WHC) and 6.2-7.1 mM NO_3_^−^ in soil moisture. Thus, at the onset of the incubation, the total amount of NO_3_^−^ per vial was 37 or 26 μ mol NO_3_^−^ in Soil3.8 or Soil6.8, respectively (see also [28]. The vials were immediately made anoxic by 6 cycles of gas evacuation and He filling [38], and incubated at 15 °C. Gases (CO_2_, O_2_, NO, N_2_O and N_2_) were measured in headspace every three hours using an autosampler linked to a GC and NO analyser [48]. At each gas sampling time point, one replicate vial of each soil type was sacrificed and NO_2_^−^ was extracted and concentration determined as described in [28]. A portion of soil from the same vial was snap frozen in liquid nitrogen then stored at −80 °C until nucleic acid extraction. The soil incubation experiment was described in detail in [28], which also explains how the rates of the four steps of denitrification were calculated based on measured concentrations of NO_2_^−^, NO, N_2_O and N_2_ throughout the incubation, and corrected for abiotic decomposition of NO_2_^−^ in the acid soil. The present MG and MT analyses were done on two of the three soils investigated by [28], and our presentation of the kinetics is limited to the calculated rates of the four steps of denitrification.

### Nucleic acid extraction

DNA and RNA were extracted from frozen samples using the method of [43] who also tested several commercial kits for these soils without success. Briefly, 3 × 0.2 g of soil was taken at time 0 (at the start of anoxic incubation) for DNA extraction, and at selected time points (0.5-27 h) during anoxic incubation for RNA extraction. Lysis was performed with glass beads in CTAB extraction buffer and phenol-chloroform-isoamyl alcohol (25:24:1), using a FastPrep-24 instrument. After ethanol precipitation, the nucleic acids were resuspended in DEPC-treated nuclease-free water purified with the OneStep PCR Inhibitor Removal Kit (Zymo Research, Irvine, USA), then split into a fraction for DNA and one for RNA. The DNA fraction was further purified using the Genomic DNA Clean & Concentrator kit (Zymo Research), then kept at −20 °C until use. The RNA fraction was digested using TURBO DNA-*free* DNase kit (Ambion, Life Technologies) according to the manufacturer’s instructions, then purified using the RNA Clean & Concentrator-5 kit (Zymo Research). Quantitative PCR (qPCR) using primers targeting the 16S rRNA gene (described below) was used to assess the presence of residual genomic DNA (gDNA) in the purified RNA fractions (defined by signal detected in the qPCR at ≤ 35 cycles), and only RNA fractions free of gDNA was used for further analysis. The purified and DNA-free RNA fractions were reverse transcribed using the Maxima Reverse Transcriptase with random hexamer primers (Thermo Scientific), according to the manufacturer’s instructions. Primers targeting the 16S rRNA or *nosZ* genes (described below) were used in qPCR to assess the quality (defined by uninhibited amplifiability) of purified DNA and reverse-transcribed cDNA.

### Sequencing the metagenome (MG), metatranscriptome (MT), and 16S rRNA genes

Triplicate DNA and duplicate RNA samples were sent for metagenomic and metatranscriptomic sequencing at The Roy J. Carver Biotechnology Center (CBC)/W. M. Keck Center for Comparative and Functional Genomics at the University of Illinois at Urbana-Champaign, using HiSeq 2500 technology. All nucleic acids were shipped in a liquid nitrogen vapour dry shipper (Cryoport) and arrived within 5 days (the Cryoport Express dewar is able to maintain the temperature at −150 °C during shipment for 10 days). The RNA integrity (including confirmation of the absence of gDNA) was also independently verified by the CBC prior to sequencing the samples. DNA samples were sent for 16S rRNA community analysis using Illumina MiSeq technology (2 × 300 bp paired-end sequencing with V3 chemistry) at StarSEQ GmbH, Mainz (Germany). The primers used targeted the V4 region of the 16S, 515f and 806rB [49, 50], as detailed by the Earth Microbiome project (http://www.earthmicrobiome.org/emp-standard-protocols/16s/).

### 16S rRNA amplicon sequence analysis

Processing of the sequenced 16S rRNA amplicons was performed by StarSEQ Gmbh, Mainz (Germany). Briefly, the sequences were demultiplexed and the adapters were trimmed locally on the MiSeq instrument with the Illumina Metagenomics 16S rRNA application, using default settings. Amplicon sequence data was processed using the Greenfield Hybrid Analysis Pipeline (GHAP) which combines amplicon clustering and classification tools from USearch [51], and RDP [52], combined with custom scripts for demultiplexing and OTU table generation [53]. Reads were subject to quality trimming with a cut-off threshold of 25 and length trimming where only reads in the length range 250 to 258 bp were retained. Trimmed forward and reverse reads were then merged and only successfully merged reads retained. OTU clustering was performed at a 97% sequence similarity. Representative sequences from each OTU were classified by finding their closest match in a set of reference 16S sequences, and by using the RDP Naïve Bayesian Classifier. The RDP 16S training set and the RefSeq 16S reference sequence collection were used for classification. NumPy, SciPy, scikit-learn and Matplotlib were used for calculation and plotting of principle component analyses (PCAs) [54–56]. Samples were rarefied to 38287 reads/sample for the purpose of determining species richness and Peilou’s evenness, calculated using USearch v11 [57].

### Analysis of MG- and MT sequences

After trimming the number of reads (in millions) was between 25.9-31.4 in the MG samples and between 16.4-42.0 in the MT samples except for two samples which were lower (4.0 for one of the 12h duplicates from Soil6.8 and 8.5 for one of the 0.5h duplicates from Soil3.8). The sequenced Illumina HiSeq reads were quality controlled using BBDuk from the BBTools package version 35.66 (http://sourceforge.net/projects/bbmap/). For functional annotation, reads were aligned using DIAMOND with an e-value cutoff of 1 × 10^−3^ [58]. For analysis of reads derived from genes involved in nitrogen metabolism comparison was done against a custom dataset described in [59, 60]. The DIAMOND output was converted to m8 blast format and analysed in R. Reads must have had a matching region of >30 amino acids and an identity of >60% to be considered matching. Output of matching reads were normalised to reads per million of total reads, RPM (see below).

Reads derived from specific genes and meeting the assigned quality cutoffs were extracted from read sets using filterbyname from the BBTools suite of programs. The extracted reads were then uploaded to KBase [61] and taxonomic assignment was performed using KAIJU using default settings [62].

### Statistical and quantitative analysis of meta-omic data

All reads counts were normalised for sequencing depth, generating RPM values: (number of reads)/(total reads that passed quality control)×10^6^. All statistical analyses and graphing were performed using in-house R scripts custom created for this purpose.

### Quantitative amplification-based analysis

The genes encoding 16S rRNA and the three denitrification reductases *nirK*, *nirS* and *nosZ* clade I were quantified by qPCR using the primers 27F and 518R for the 16S rRNA gene [63], 517F and 1055R for the *nirK* gene [64], cd3aF and R3cd for the *nirS* gene [65] and Z-F and 1622R for the *nosZ* gene [66]. DNA samples were diluted to 1-10 ng of DNA per reaction. All cDNA and RNA samples (DNase-digested) were used without dilution. Each 20 μL qPCR reaction contained SYBR *Premix Ex Taq* II (Tli RNaseH Plus) (Takara Bio) and was run according to the manufacturer’s instructions and included 0.4 μM of each primer and 2 μ L of template. The optimised qPCR cycling conditions for all primer sets were 95 °C for 5 min, 40 cycles of 95 °C for 30 s, *x* for 60 s, 72 °C for 30 s, 82 °C for 20 s, and a final melting curve analysis from 60 °C to 95 °C to determine the specificity of amplicons, where *x* = 54 °C (16S rRNA gene), or 60 °C (denitrification genes). To reduce background signals from primer dimers and unspecific PCR products, the fluorescence signal was measured during the final step of each cycle, at 82 °C. The detection limit of each qPCR run was 5 copies per microliter of reaction [43], which was approximately 4 × 10^2^ copies g^−1^ soil (ww).

## Results

### Kinetics of denitrification intermediates depict a pH-dependent response to anoxia

The denitrification kinetics of the two soils during 45 h of anoxic incubation are shown in Fig. 1, in which the sampling occasions for MT analyses are also indicated. A more complete description of the incubation experiment is given by [28], including detailed analyses of production/reduction rates of the denitrification intermediates/end products over 70 h. The analysis included a careful mineral N budget analysis demonstrating 100% recovery of NO_3_^−^-N as N_2_ for the soil with pH=6.8, which suggests negligible reduction of nitrate to ammonium in this soil. For Soil 3.8, the recovery as N-gas (N_2_ + N_2_O +NO) was lower (77%), but abiotic nitrosylation of organic material accounted for 17% (see Table 2 in [28], thus in total 94% was accounted for, leaving only 6% of the NO_3_^−^-loss that could possibly be ascribed to DNRA.

The two soils showed striking differences in their accumulation of NO_2_^−^ and N_2_O, while they had very similar NO kinetics (Fig. 1A). The NO_2_^−^ concentration in Soil3.8 was 20-50 μM in the soil moisture during the entire anoxic incubation, except for the 36-40 h period when it reached ~100 μM. This corroborates the general notion that NO_2_^−^ concentrations are low in acidic soils. In Soil6.8, on the other hand, NO_2_^−^ steadily increased from the beginning, reaching 2-3 mM in the soil moisture at 20 h, after which levels decreased and were undetectable at the end of the sampling period. Both soils had a transient accumulation of 3-4 μmol NO vial^−1^, which is equivalent to 1.5-2 μM NO in the soil moisture. The soils had profoundly different N_2_O and N_2_ kinetics. Soil3.8 accumulated N_2_O but little or no N_2_ during the first 35 h, while Soil6.8 accumulated both N_2_O and N_2_ at similar rates from the very beginning. The gas kinetics is reflected in the calculated rates of the four steps of denitrification (Fig. 1 B): In the acid soil, *V*_*NAR*_/*N*_*AP*_ (NO_3_^−^→NO_2_^−^), VNIR (NO_2_^−^→NO) and *V*_*NOR*_ (NO→N_2_O) were similar, while *V*_*NOS*_ (N_2_O →N_2_) was close to zero during the first 35 h. In the soil with near neutral pH, *V*_*NAR*_/*N*_*AP*_ exceeded the other enzyme rates during the first 15 h but declined rapidly to zero as NO_3_^−^ was depleted. In contrast to the acidic soil, *V*_*NOS*_ was high from the very beginning of the incubation in Soil6.8.

Based on the total NO_2_^−^-N (TNN=NO_2_^−^+HNO_2_), we calculated the concentration of undissociated HNO_2_ (aq) in the soil matrix using the Henderson-Hasselbalch approximation (see Supplementary material p. 3), which forms an equilibrium with the gas HONO in the atmosphere, thus predicting the potential emission of HONO. Despite the high accumulation of TNN in Soil6.8 (up to 3.6 mM), the concentration of undissociated HNO_2_ was ≤ 1.4 μM. In Soil3.8 the concentration HNO_2_ was almost two orders of magnitude lower (Fig. 2).

**Figure 2.**
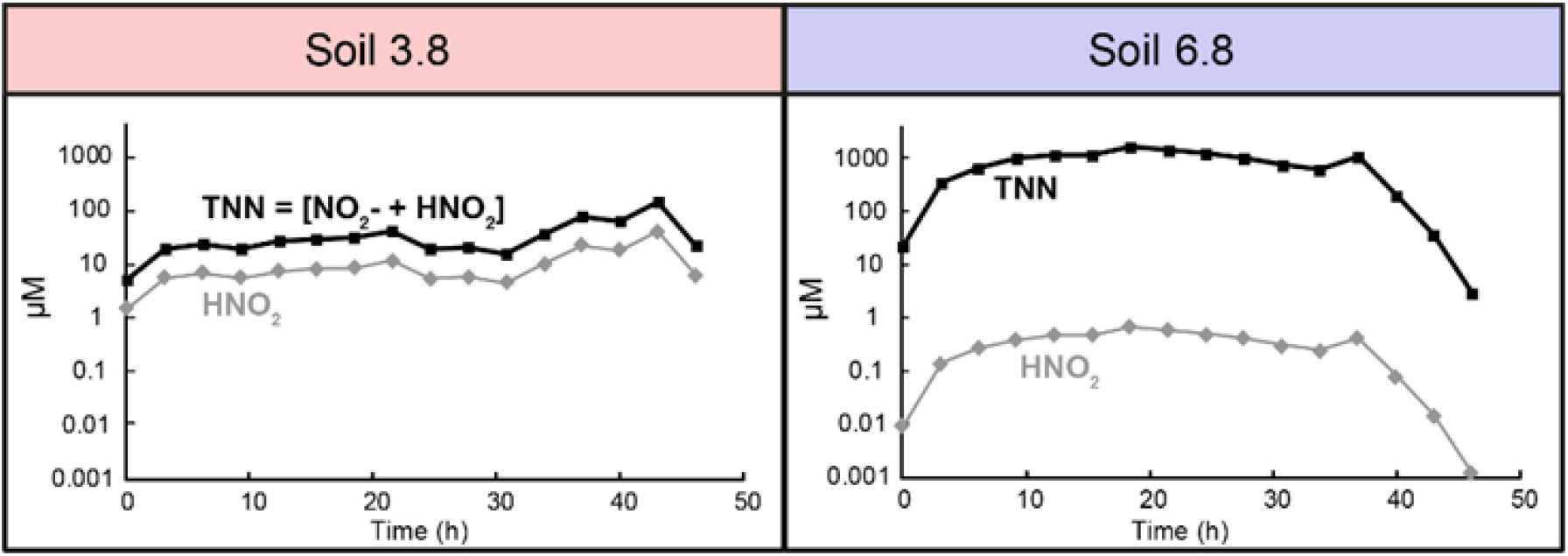
Concentrations of total NO_2_^−^-N (TNN) and the calculated undissociated HNO_2_ (μ M in soil moisture), assuming equilibrium: [HNO_2_]/([HNO_2_]+[NO_2_^−^])=1/(1+10^pH−pKa^), where p*K*_a_= 3.398.

### Soil bacterial community composition differed by pH but was stable during incubation

In the 16S rRNA gene amplicon analysis >99.29 % of all sequenced reads were annotated as bacterial, about 0.004 % were unclassified, and the rest belonged to Archaea, which represented ≤0.60 % of the reads in Soil6.8 and ≤1.03 % in Soil3.8. Principal component analysis of the bacterial 16S rRNA gene reads (Fig. S1A) clearly separated the reads from the two soils along PC1, which explained 94% of the total variation and also showed that the variation between replicate samples was low. The most abundant classified phyla in both soils were Proteobacteria, Acidobacteria, Actinobacteria, Planctomycetes, Verrucomicrobia and acteroidetes. A breakdown of Proteobacteria showed that Alphaproteobacteria were most abundant in both soils, followed by Beta-, Gamma-and Deltaproteobacteria (Fig. S1B). The main differences between the soils was a larger relative abundance of Actinobacteria and Alpha- and Betaproteobacteria in Soil3.8 than in Soil6.8 along with a higher relative abundance of Bacteroidetes in Soil6.8 than in Soil3.8. The OTU richness was higher in the Soil6.8 samples with 2846 OTUs observed on average for the Soil6.8 samples compared to an average of 1882 OTUs for the Soil3.8 samples. Soil6.8 samples were also slightly but significantly (p=1.66×10^−5^) more even according to Peilou’s evenness measure with Soil6.8 samples having an average Peilou’s evenness measure of 0.814 and compared to 0.796 for Soil3.8 [57]. The microbial community profile of Soil6.8 was stable during the 27 h incubation, suggesting that differences observed in the MT can be reasonably attributed to variations in transcription patterns, and not due to bacterial growth causing a shift in the bacterial community composition.

### Occurrence and prevalence of reads in the MG related to denitrification and DNRA

Reads annotated as NAR, here defined as *nap*+*nar* reads, were twice as abundant as NIR, NOR and NOS gene reads in the MG of both soils (Fig. 3A; Table 1). Of the two types of NAR, *narG* reads were 6.1±0.3 times more abundant than *nap* reads in Soil3.8 but only 2.1±0.3 times more abundant in Soil6.8 (Table 1). Levels of NIR (*nirK*+*nirS*) were comparable in the two soils with RPM values of 36.4±1.7 and 44.6±1.4 for Soil3.8 and Soil6.8, respectively (Table 1). *nirK* genes were much more abundant than *nirS* in both soils with a *nirK/nirS* ratio of about 40 in Soil3.8 and 7 in Soil6.8. The NOR gene reads (*cnor*+*qnor*) were more abundant in Soil3.8, where RPM values were 74.9±0.5 compared to 48.1±0.3 for Soil6.8. The *qnor* genes dominated over *cnor* in both soils and were about 11 and 4 times higher in Soil3.8 and Soil6.8, respectively. Total NOS reads (*nosZ* clade I+II) were instead somewhat higher in Soil6.8 than in Soil3.8 (25.8±2.0 *vs* 18.2±1.0). To summarize this for all genes, the order of gene reads from highest to lowest abundance was for Soil3.8 *narG* > *qnor* > *nirK* > *napA* > *nosZ* clade I > *cnor* > *nirS >nosZ* clade II; and for Soil6.8 *narG* > *napA* > *nirK* + *qnor* > *nosZ* clade II > *cnor* > *nirS* > *nosZ* clade I.

**Figure 3.**
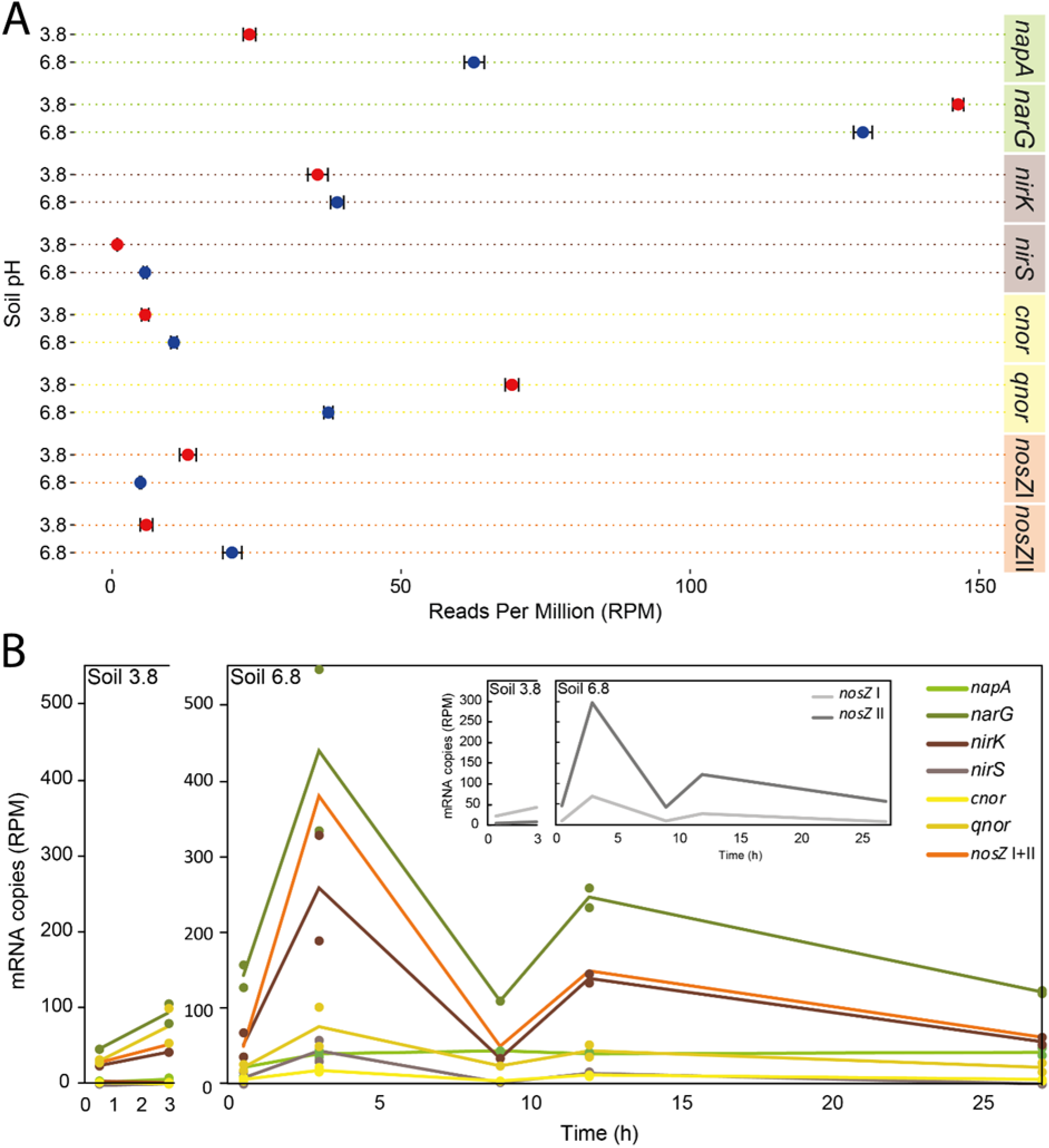
Genetic potential and transcription of selected denitrification genes. **A.** Metagenome analysis showing a comparison of the abundance of denitrification genes in Soil3.8 and Soil6.8 based on number of reads annotated to the gene in question per total number of reads in the sample. Averages of triplicate soil samples taken at the start of the incubation for metagenomics analysis. Bars indicate standard deviation. B. Metatranscriptome analysis showing changes in the abundance of denitrification gene transcripts during the first 3 hours of incubation for soil3.8 and 27h for soil6.8. Averages (lines) and single data points are shown for duplicate soil samples taken after 0.5, 3, 9, 12 and 27 h of anaerobic incubation. Reads of *nirS* transcripts were not detected in Soil3.8. The insert shows the transcriptional dynamics of two *nosZ* clades only, for clarity.

**Table 1.**
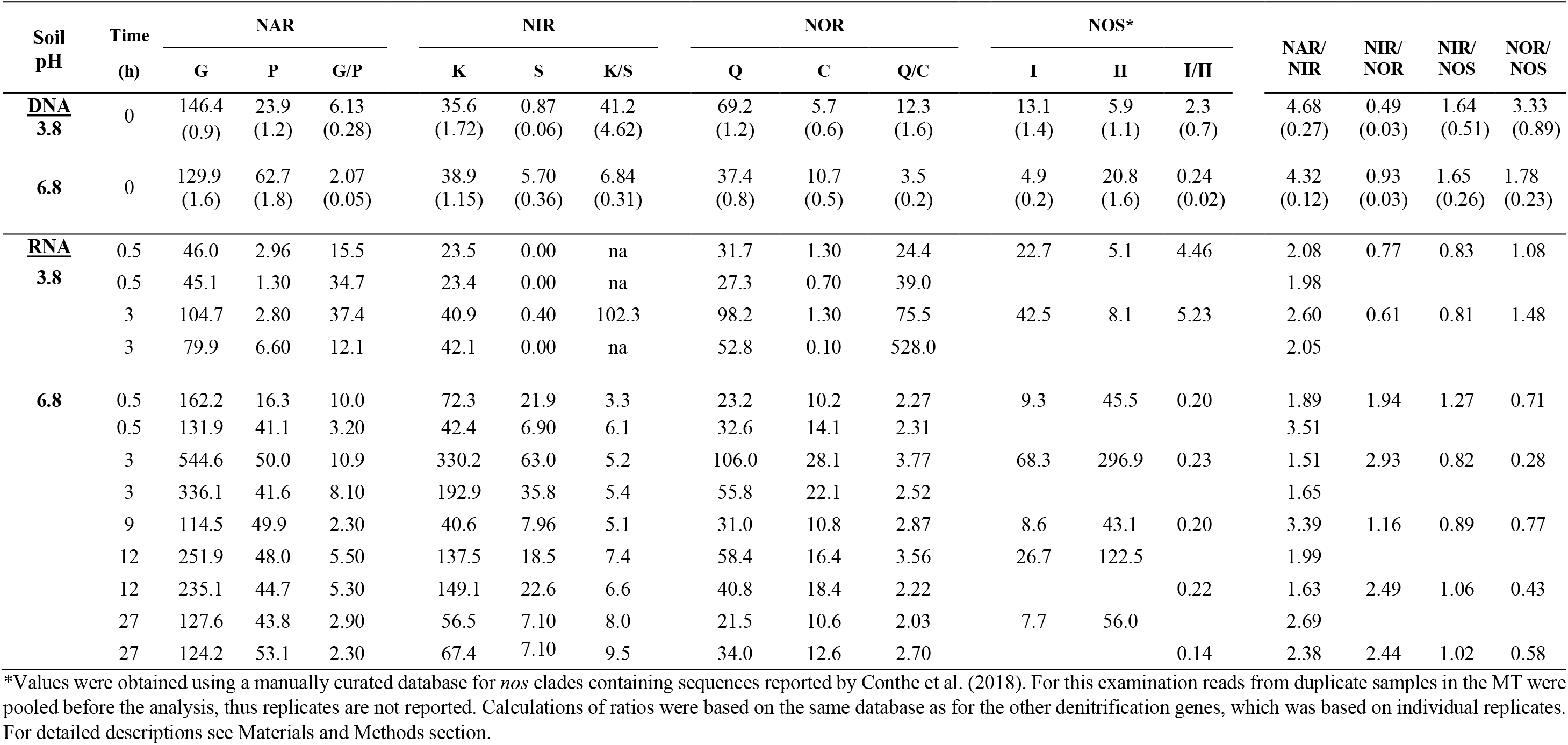
Abundances of gene and transcript reads in the metagenome (DNA) and metatranscriptome (RNA) of two soils with pH 3.8 and 6.8 during anoxic incubation with NO_3_^−^. NAR=nitrate reductases, sum of *narG* and *napA* abundancies; NIR=nitrites reductase, sum of *nirK* and *nirS* abundancies; NOR = nitric oxide reductases, sum of *qnor* and *cnor* abundancies; NOS= nitrous oxide reductases; sum of clade I and clade II abundancies. DNA samples were taken just before the start of the incubation (t=0 h, Fig. 1A). RNA samples were taken after 0.5 and 3 h incubation (Soil 3.8) and after 0.5-27 h incubation (Soil 6.8) (Fig. 1A). All values are in reads per million (RPM). Ratios of reads from selected genes/transcripts are also given. DNA: Average values are given for abundance of individual genes, values in parenthesis are standard deviation; n=3. RNA: Duplicate metatranscriptome samples were analyzed from each sampling time except for the 9 h sample from Soil6.8 for which one samples was lost. Individual values for read abundances and ratios are shown except for the *nosZ* clades for which data from duplicate samples were pooled. Ratios including NOS are therefore calculated from average values. na= not applicable.

Since the two soils accumulated different amounts of denitrification intermediates, we calculated ratios of MG reads representing the four steps of denitrification (Table 1), to examine if the genetic potential for production/consumption of the different intermediates was related to the net production seen in Fig. 1. The NAR/NIR ratios were similar in the two soils (4.68±0.27 for Soil3.8 and 4.32±0.12 for Soil6.8), which, by itself, is not consistent with the higher net production of NO_2_^−^ in Soil6.8. The NOR genes were more abundant than NIR in Soil3.8, with a NIR/NOR ratio of 0.49 ±0.03 compared to 0.92 ± 0.03 in Soil6.8. NIR and NOR were more abundant than NOS in both soils with NIR/NOS ratios of 2.83 and 2.07 and NOR/NOS ratios of 5.81 and 2.23 in Soil3.8 and Soil6.8, respectively. The two *nosZ* clades (I and II) differed in gene abundance in the two soils with higher values for clade I in Soil3.8 and *vice versa* in Soil6.8 (Fig. 3A). The *nosZ* clade I/clade II ratio (based on RPM values) was 28.1 in Soil3.8 and 0.25 in Soil6.8 (Table 1).

We also analysed accessory genes of the *nos* operon, with a focus on clade I, using a manually curated database. The *nosR* gene is a part of the *nos* operon in clade I but not in *nosZ* clade II organisms [15, 45]. Its product is essential for N_2_O reduction in clade I organisms and is suggested to be involved in transcription of *nosZ*, and also in electron transfer to the NosZ reductase [46]. In accordance with *nosZ* clade I being dominant in Soil3.8, we found about twice as many reads derived from *nosR* in Soil3.8 than in Soil6.8 with RPM values of 16.2±0.3 *vs* 7.0 ±0.3, respectively. Moreover, we examined the relative abundance of the accessory genes *nosL, D* and *Y*, which are found both in *nosZ* cladeI and clade II organisms [15, 45]. For each of these genes, the read abundance was similar in the two soils (Fig. 4), which is in accordance with the comparable abundance of NOS (*nosZ* clade I+II) in the two soils (12.5 RPM in Soil3.8 and 20.8 in Soil6.8, Table 1). The gene reads for *nosF*, which is also part of the *nos* operon both in clade I and II [45], were 10-45 times higher than for the other accessory genes which points to uncertainties in the databases for this gene.

**Figure 4.**
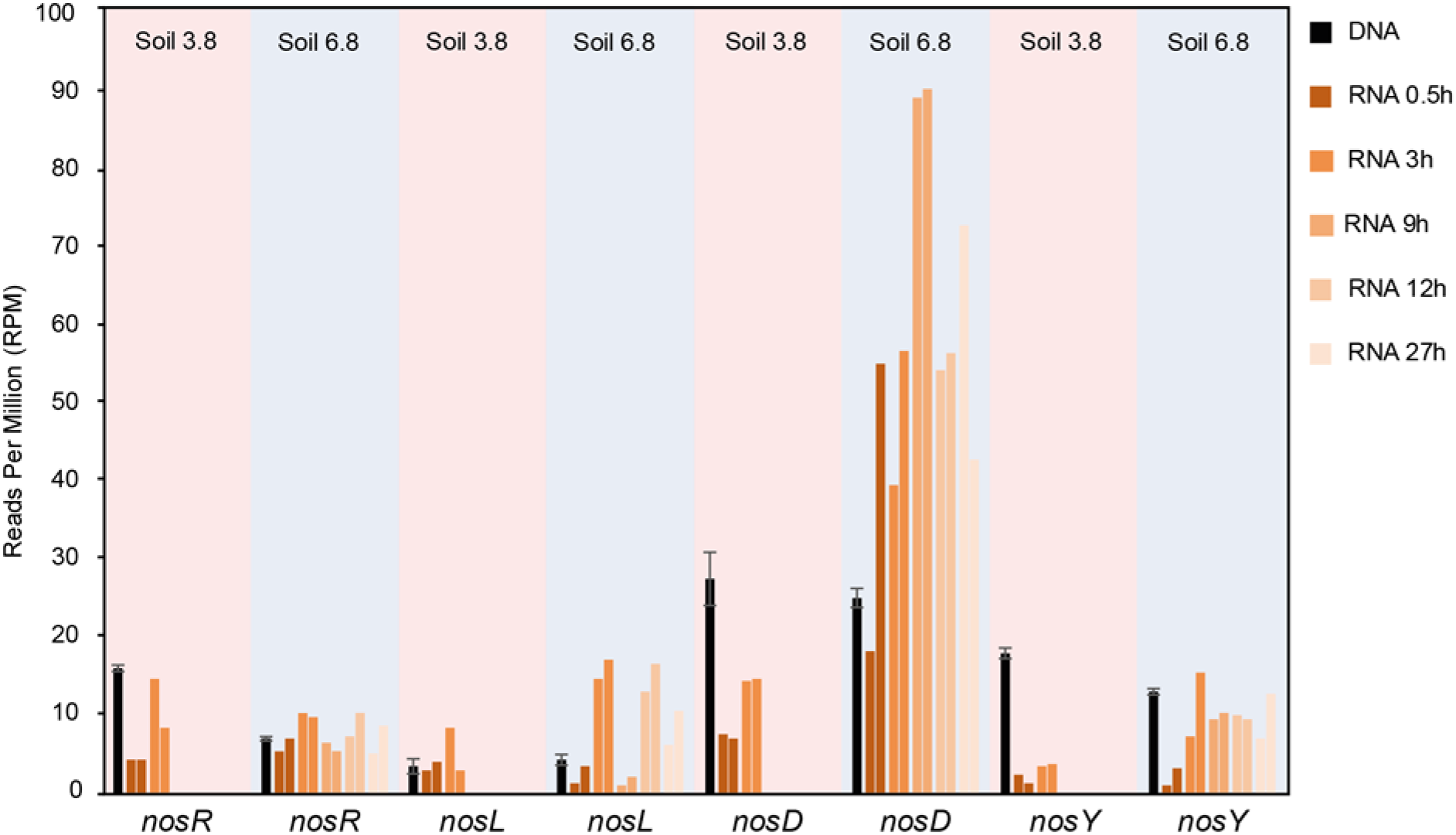
Genetic potential and transcription of selected genes in the *nos* operon in the metagenome and metatranscriptome of Soil3.8 and Soil 6.8. Black bars show average gene read abundances in the metagenome (n=3; bars show sd). Colored bars show transcript read abundances after 0.5 and 3h (Soil3.8) and after 0.5, 3, 9, 12 and 27 h of incubation (Soil6.8). Duplicate samples were analyzed for each sampling point, shown as individual bars with the same color.

In addition to the canonical denitrification genes, we examined reads from the two DNRA-related genes *nrfA* and *nirB*. They were about 3 times more abundant in Soil6.8 than in Soil3.8 with *nirB* dominating over *nrfA* in both soils (Fig. S2). We also examined the occurrence of reads derived from the gene *hmp*. This gene, reported from a wide range of bacteria [67], encodes a flavohemoprotein that has a role in NO detoxification. Unlike the NOR genes, we found similar and relatively low abundances of *hmp* in the two soils (16±0.8 and 18±1.4 RPM, not shown).

### MT reads related to denitrification and DNRA

Earlier PCR-based investigations of soils from the same site as that studied here showed that denitrification genes were quickly transcribed upon onset of anoxia [24, 38]. Based on these findings, RNA samples for the present study were taken from both soils 0.5 and 3 h after the start of the experiment (red arrows, Fig. 1) to characterize the community response to denitrifying conditions. Additional RNA samples were taken from Soil6.8 prior to the peaks in NO_2_^−^, NO, and N_2_O (Fig. 1) to determine if there were new bursts of gene transcription during the incubation and, if so, if these were from already active organisms, or if different organisms became transcriptionally active at the later time points. The transcript read abundancies are shown in Fig. 3B. For clarity, the *nosZ* genes are shown as the sum of the two clades in the main figure, while the insert shows the clades separately. The reads in the meta-transcriptome from Soil6.8 exhibited a strong increase between 0.5 and 3 h for almost all denitrification gene transcripts. Over the same time frame there was a smaller, yet substantial increases for several of the genes in Soil3.8. Two distinct spikes in denitrification gene transcription were seen at 3 and 12 h, particularly for *narG*, *nirK*, *qnor* and *nosZ* (Fig. 3B). Reads of *narG* represented the most common denitrification transcripts in both soils and at all time points. The overall trend in transcript read abundance in Soil6.8 was *narG* > *nosZ* > *nirK* > *qnor* > *napA* > *nirS* > *cnor*, almost without exception, throughout the incubation (Table 1). The trend for Soil3.8 was *narG* > *qnor* > *nosZ* > *nirK* > *napA* > *cnor* = *nirS*, the main difference compared to Soil6.8 being that *qnor* was second most abundant, followed by *nosZ*. Similar to the gene reads in the MG, *nosZ* clade I transcript reads were more abundant than clade II reads in Soil3.8 and vice versa for Soil6.8.

It could be argued that the ratios of transcripts representing the reductases responsible for the production and consumption of the various denitrification intermediates are more suitable to use for comparison with phenotypic data than actual RPM values (transcript read abundances and selected ratios are given in Table 1). The NAR/NIR ratios were very similar in the two soils with average values of 2.2±0.3 for Soil3.8 and 2.3±0.8 for Soil6.8, with some variations between sampling times. The NIR/NOR ratios, on the other hand, were almost 3 times higher in Soil6.8 compared to Soil3.8 (average values including all sampling times were 2.2±0.8 and 0.7±0.2, respectively). The high NOR transcription in Soil3.8 most likely contributed to keeping NO emissions low, despite known chemical reactions between NO_2_^−^ and soil compounds producing NO. The NIR/NOR ratios of Soil6.8 for the individual sampling points further showed that NOR was consistently lower than NIR throughout the 27h incubation with NIR/NOR ratios of 1.2-2.5. Transcription of NOS (sum of clade I, clade II and ambiguous reads) was higher in Soil6.8 than in Soil3.8 and read numbers increased almost 7 times from 56 to 380 in Soil6.8 between 0.5 and 3 h. In Soil3.8 the NOS transcript reads almost doubled during the same period. NIR/NOS ratios were rather similar in the two soils, around 0.8 in Soil3.8 and 1.0 ±0.2 in Soil6.8. NOR/NOS ratios were higher in Soil3.8 than in Soil6.8, with an average of 1.3±0.3 for the first two sampling points while, during the same time, this ratio for Soil6.8 was 0.6±0.2. These ratios reflect the higher number of NOR reads in Soil3.8, combined with higher reads for NOS in Soil6.8.

Transcript reads of *nos* accessory genes were detected in both soils at all sampling occasions (Fig. 4). The transcriptional activity increased for all these genes between 0.5 and 3 h, which is a strong indication that the lack of N_2_O reduction in Soil3.8 was not caused by any transcriptional control mechanism. The *nosR* transcript abundances were comparable in the two soils.

We also examined the MT with regard to the two DNRA genes *nfrA* and *nirB,* expecting low read abundance since no DNRA activity was discerned from the gas analyses. Surprisingly, these two genes were transcribed at levels comparable to the NIR and NOR genes, with sometimes high read values for *nirB* (Fig. S2).

### PCR-based quantification of functional genes and transcripts overlooked substantial parts of the community

The abundances of some of the denitrification genes and transcripts in the MGs and MTs (Fig. 3; Table 1) were compared to qPCR-results in samples from the same soil incubation (Fig. 5), although more time points were included in the qPCR analysis. We targeted *nirK*, *nirS* and *nosZ* clade I using standard primer pairs (see Materials and Methods). The comparison revealed some striking differences between -;omics and PCR-based results. While the “-omics” based results showed dominance of *nirK* over *nirS* both in the MGs and MTs, the qPCR-based quantifications showed 1-2 orders of magnitude lower abundances of *nirK* genes and transcripts compared to *nirS,* except for Soil3.8 where the difference was smaller. Comparing the *nirS* and *nosZ* (clade I) abundancies shows nearly identical abundancies of these genes in Soil6.8 according to the MG (Fig. 3A) while qPCR showed almost 10 times fewer *nosZ* than *nirS* gene copies based on qPCR (Fig. 5A). The MG- and qPCR-based results from Soil3.8 showed better correspondence, with *nosZ* being more abundant than *nirS*. For the transcripts (Figs. 3B and 5B), *nosZ* was more abundant than *nirS* in Soil3.8 according to the MT analysis, while the qPCR showed an opposite trend. Similarly, qPCR gave *nosZ* transcript levels that were 5-10 times lower than *nirS* in Soil6.8, while their levels were similar in the MT. The results indicate a critical primer bias in the amplicon-based quantification leading to severe underestimations of *nirK* abundance and overestimations of *nirS* compared to *nirK* and *nosZ* clade I.

**Figure 5.**
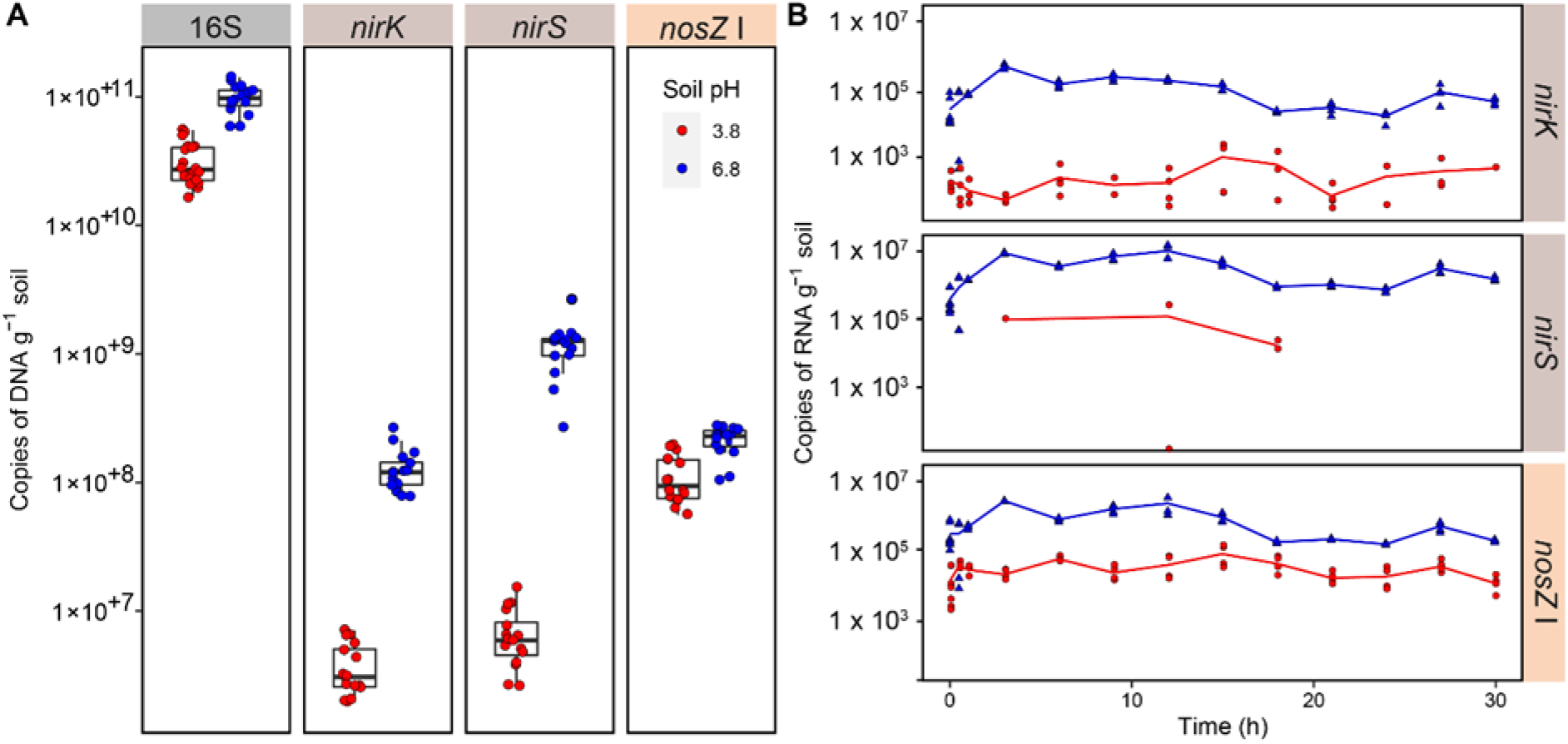
Amplification-based quantification (qPCR) of genes and transcripts (copies g^−1^ soil, wet weight). (A) Quantification of gene copies. Primers targeting the 16S rRNA (27F/518R), *nirK* (517F/1055R), *nirS* (cd3aF/R3cd) and *nosZ cladeI* (nosZF/1622R) genes were able to detect gene copies in both soils (B) mRNA transcripts of *nirK*, *nirS*, and *nosZ*. Red= Soil3.8; Blue= Soil6.8.

### Taxonomic annotation of denitrification genes and transcripts

Each gene and corresponding transcript had a unique taxonomic profile that varied by soil pH (Fig. 6). Proteobacteria, Actinobacteria and Bacteroidetes were the most abundant phyla of denitrifiers in the MG and MT in both soils. Reads assigned to these phyla were detected for most denitrification genes and transcripts with the exceptions of Bacteroidetes, which were not found among *narG* and *nirS* reads, and Actinobacteria, which were not found among the *nirS*, *cnor* and *nosZ* clades I and II reads. Several other phyla such as Firmicutes, Chlamydiae, Nitrospira, Spirochaetes and Verrucomicrobia were represented by reads only from one or a few genes/transcripts.

**Figure 6.**
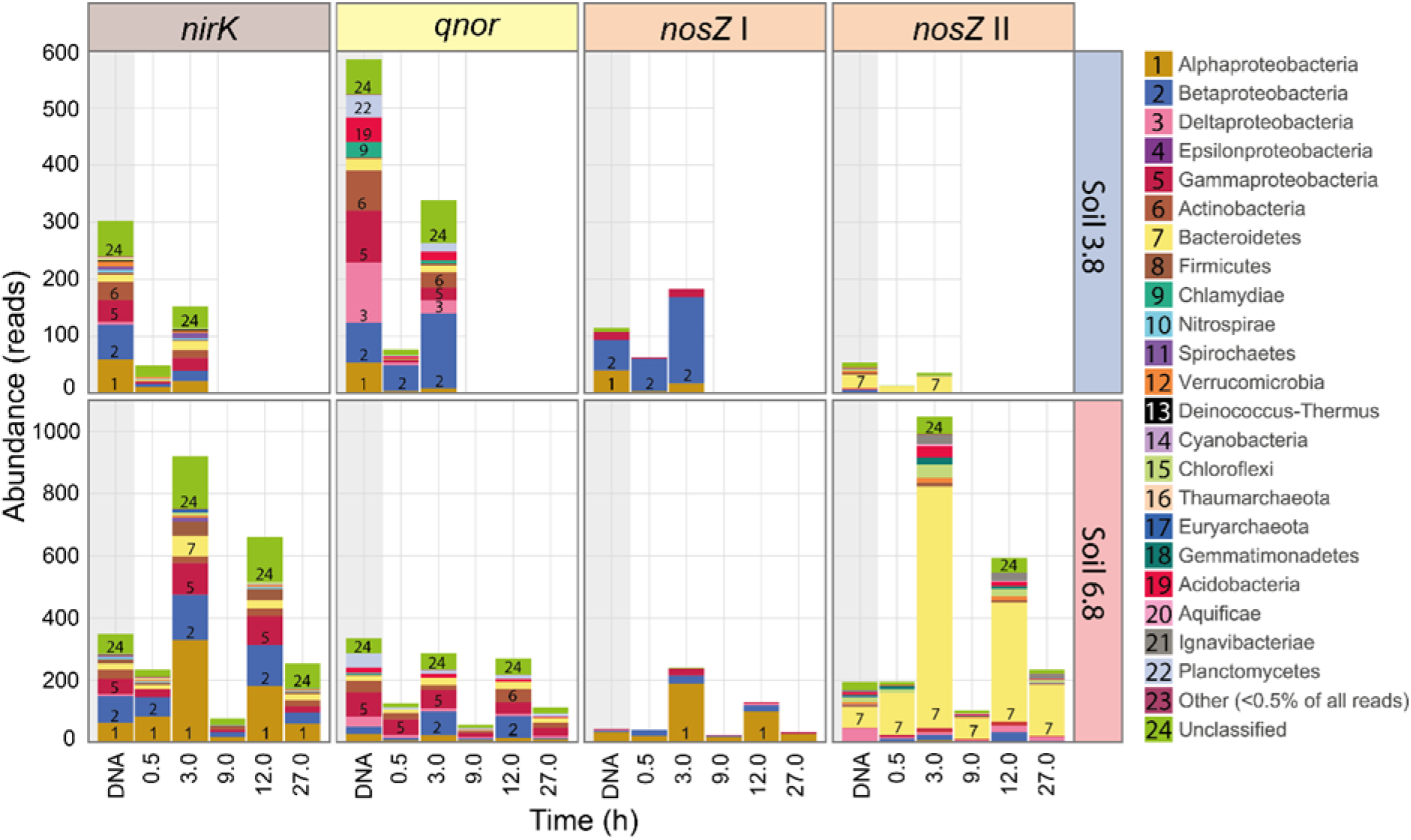
Denitrification gene and transcript prevalence as summed reads across replicate samples (n=3 for MG; N=2 for MT). Annotated genes (MG; DNA sampled at start of incubation) and transcripts (MT) sampled after 0.5 and 3 h anoxic incubation (Soil3.8) and 0.5-27 h anoxic incubation (Soil6.8) (for sampling points also see Fig. 1A).

The MG reads of *narG*, the most abundant among the denitrification genes, were dominated by Proteobacteria, mostly the classes Alpha- and Betaproteobacteria, and Actinobacteria (Table S1). Second most abundant in both soils were *napA* reads, which were mainly derived from Proteobacteria. Organisms belonging to these phyla also dominated the transcriptional activity for *narG* and *napA* genes. *narG* reads from Nitrospira were high as well in Soil6.8. MG reads derived from the NO_2_^−^ reduction gene *nirK*, which was far more abundant than *nirS* in both soils (Table 1), were attributed to a number of phyla which were dominated by Proteobacteria, primarily Alpha-Beta- and Gamma proteobacteria, and by Actinobacteria, Bacteriodetes and Firmicutes. These same phyla also dominated the MT reads of *nirK* in both soils, with Proteobacteria showing particularly high transcription of this gene in Soil6.8 at the 3h sampling time. MG reads from *nirS* were predominantly from Betaproteobacteria in Soil6.8, which also accounted for the highest *nirS* transcriptional activity. Only 7 classified *nirS* MG reads were identified in Soil3.8 compared to 242 *nirK* reads (Table S1). Similar as for the *nirK* and *nirS* genes, the *qnor* genes dominated strongly over *cnor* genes in the MG, and the *qnor* genes belonged to several phyla while the *cnor* reads were mostly from Proteobacteria. The Delta- and Gammaproteobacteria had the highest number of *qnor* MG reads in both soils, but it was the Betaproteobacteria that dominated among the transcripts showing comparatively high and immediate transcription in Soil3.8. Interestingly, *qnor* MG and MT reads from Acidobacteria and Planctomycetes were detected in both soils at relatively high abundance. Apart from a few acidobacterial *nosZ* clade II reads, reads from these two phyla were not found for the other denitrification genes in the MG or MT.

The total number of *nosZ* MG reads was higher in Soil6.8 than in Soil3.8 (212 *vs* 152, Table S1). As expected, *nosZ* clade I MG and MT reads were only from Proteobacteria in both soils. *nosZ* clade II MG reads were more diverse, especially in Soil6.8, and were comprised of not only Proteobacteria, mostly Deltaproteobacteria, but also several other phyla including Bacterioidetes, which was the most highly represented group, as well as Firmicutes, Verrucomicrobia, Chloroflexi, Gemmatimonadetes and Acidobacteria among others. The transcriptional activity of *nosZ* clade I was dominated by Alphaproteobacteria in Soil6.8 and by Betaproteobacteria in Soil3.8. The pattern was very different for transcription of *nosZ* clade II, which was dominated by Bacteroidetes in both soils. The transcript abundance of clade II in Soil3.8 was low but increased between 0.5 and 3 h and included not only Bacteriodetes but also a few reads from Proteobacteria, Bacteriodetes, Chloroflexi, Gemmatimonadetes, Ignavibacteriae and Acidobacteria. Taken together, the results for the two *nosZ* clades show that the problem of producing functional NosZ in Soil3.8 was common to several phyla. Soil6.8 showed high transcriptional activity of *nosZ* clade II, dominated by Bacteriodetes, but transcripts were also detected from all other phyla for which *nosZ* clade II were registered in this soil, except Euarchaeota.

## Discussion

Liming, which has traditionally been a means to improve the fertility of acidic soils [68], is known to cause changes in microbial community composition and diversity [69, 70]. This was also seen in the present study, where the PCA of the 16S rRNA gene sequences clearly separated the two soils (Fig. S1A). The sequence analysis also revealed a slight but significant increase in diversity in the limed soil. The dominant phyla in both soils were those that generally dominate in soil studies [71] and no major differences between the soils were seen on the phylum level, or on class level for the proteobacteria, meaning that changes in the abundance of different groups had taken place at lower taxonomic levels. During the 27 h anoxic incubation, when Soil6.8 was monitored with metatranscriptomics, the community of Soil6.8 remained stable as judged from the 16S rRNA gene analysis (Fig. S1B), suggesting that any differences in the MT can be reasonably attributed to transcriptional regulation patterns and not caused by growth of some populations.

The accumulation of NO_2_^−^ and N_2_O in the two soils shows contrasting patterns, with low NO_2_^−^ and high N_2_O levels in Soil3.8 and *vice versa* for Soil6.8. The pattern of NO accumulation was more similar in these two soils, as presented in [28]. In the present study we investigated if the contrasting denitrification phenotypes of the two soils could be predicted from the abundance and transcription of the denitrification genes. Net production of denitrification intermediates will take place as soon as the reduction rate of one of the denitrification steps surpasses that of the following step. The NO_3_^−^ reduction rate (*V*_*NAR*_) in Soil6.8 grossly exceeded the NO_2_^−^ reduction rate (*V*_*NIR*_), as long as nitrate was present (Fig. 1B and [28]). The MT analysis showed higher transcription of NAR (*napA*+*narG*) than of NIR (*nirK*+*nirS*) (Fig. 3B) with NAR/NIR ratios between 1.6 and 3.4 (Table 1). The ratio was similarly high for Soil3.8, however (2.0 and 2.3 for the two time points measured in this soil), despite the near-absence of NO_2_^−^ accumulation in this soil. Thus, no direct link was found between the transcript ratio NAR/NIR and NO_2_^−^ accumulation or the *V*_*NAR*_/*V*_*NIR*_ ratio, as affected by pH. In theory, abiotic NO_2_^−^ decomposition could be the primary reason for the marginal transient accumulation of NO_2_^−^ in Soil3.8, but this was refuted by the careful analyses of Lim *et al.* [28], who reached the conclusion that enzymatic NO_2_^−^ reduction was the major sink for NO_2_^−^ in Soil 3.8. One explanation for the marginal NO_2_^−^ accumulation in Soil3.8 could be that NO_2_^−^ reductase has a low pH optimum, as shown in a study by Abraham *et al.* [72] where measured NirK activity *in vitro* was ~4 times higher at pH 4.2 than at pH 7. Thus, Soil3.8 could have a much lower V_NAR_/V_NIR_ ratio than Soil6.8, despite the nearly equal NAR/NIR-ratio for the two soils. The results demonstrate that transcript numbers or ratios of transcript numbers are poor predictors of metabolic activities in soils. Even more evident is the discrepancy between gene numbers and activity. The NAR/NIR ratio in the metagenome was almost identical in the two soils (Table 1), despite the substantial difference in NO_2_^−^ accumulation. The understanding of how denitrifiers control NO_2_^−^ levels is far from complete, and different phenotypes have been described which are probably all present in complex soil microbial communities [18]. Some organisms perform complete denitrification of NO_3_^−^ to N_2_ with little or no accumulation of intermediates, while others show complete inhibition of *nir* transcription until available NO_3_^−^ to NO_2_^−^ before further reduction takes place [23], and yet others reduce about half of the provided NO_3_^−^ to NO_2_^−^ before further reduction [73].

One reason that the NAR genes were by far more abundant than the genes for the other denitrification steps (Fig. 3A) could be that these genes are also carried by organisms performing DNRA [74]. Organisms with only NAR but lacking genes both for DNRA and denitrification are, however, also common [19], and in a study of bacterial isolates from the same field site as in the present study, 18 and 19% of the isolates from acidic and neutral soil, respectively, performed only NO_3_^−^ reduction to NO_2_^−^ without any further reduction [18]. The present study showed a strong dominance of *narG* over *napA* both in the MGs and in the MTs (Table 1; Fig. 3), despite the common occurrence of bacteria with *napA*+*narG* or with only *napA* among hitherto studied DNRA- and denitrifying organisms [75]. Still, *napA* was 2.6 times more abundant in the metagenome of Soil6.8 compared to Soil3.8 (Fig. 3A), in line with the significant positive relationship between *napA* and pH reported by [59], but the reason for this apparent pH dependence on the abundance of *napA* is not clear.

According to the MG analysis the abundance of NIR genes was similar in the two soils, with strong dominance of *nirK* over *nirS*, especially in Soil3.8 where *nirS* genes were almost absent (Table 1; Fig 3A;). This contradicts the general conception that *nirS* is more abundant in most environments, which is derived from primer-based studies [38, 76, 77]. However, while *nirS* is mainly found in Proteobactera, *nirK* is spread among taxonomically diverse groups, many of them being non-proteobacterial denitrifiers [16, 17, 78], which was also found in the present study (Fig. 6). Transcripts of *nirK* dominated over *nirS* also in the MT (Fig. 3B), and represented a diverse range of phyla including, in addition to Proteobacteria, also Actinobacteria, Bacteroidetes and Firmicutes (Fig. 6). This is in accordance with the results from a non-primer based study of another agricultural soil by [77]. The occurrence of a taxonomically diverse group of denitrifiers actively transcribing *nirK* probably reflects that this gene is easily transferred horizontally and expressed in new hosts since it does not require accessory genes to produce a functional enzyme, in contrast to *nirS*, which is part of a multi-gene operon [78]. This, and other evolutionary mechanisms, are a likely explanation to the reported incongruences between *nirK* and 16S rRNA phylogenies [16, 79]. Recently, Nadeau *et al.* [59] pointed out that denitrification is a non-essential trait for microbes, and that genes are gained and lost by individual members of the denitrifying community depending on the conditions. It is likely that this applies in particular to the functional genes *nirK* and *qnor*, which are probably more easily transferred horizontally than the other denitrification genes, and it can be speculated that they comprise a pool of genes that circulate between organisms depending on their needs.

The complete recovery of all added NO_3_^−^ as N_2_ in Soil6.8 (Fig. 1A) indicated minimal or no conversion of NO_2_^−^ to NH_4_^+^ and thus that the NO_2_^−^ produced from NAR activity was not used by DNRA organisms. This is surprising, taking into account the relatively high abundance of reads from *nrfA* and *nirB* genes and transcripts (Fig. S2). Likewise, DNRA was probably insignificant in Soil3.8, since 94 % of the NO_3_^−^ loss in this soil could be accounted for by N-gas + nitrosylation (see [28]. In theory, full recovery of NO_3_^−^ reduction as N-gas could be obtained in a system with equal rates of DNRA and anammox. However, members of *Brocardiales*, which comprises the anammox-*Planctomycetes* [80], were scarce in the 16S rRNA gene analysis (<0.008% of 16S reads in all SoilA samples), which lends little support to this hypothesis.

The net production of NO was similar in the two soils despite the substantial abiotic reduction of NO_2_^−^ that contributed to NO emissions from Soil3.8 [28], in addition to the enzymatic NO_2_^−^ reduction taking place in both soils. The strong control of NO in Soil3.8, seen from the calculated *V*_*NOR*_ activity (Fig. 1B), is in agreement with the 2.5 times increase in *qnor* transcript abundance between 0.5 and 3 h (Fig. 2B) and NIR/NOR ratios <1 (Table 1). In line with this, the gene abundance of *qnor* was also high in Soil3.8, almost 2 times its abundance in Soil6.8 (Fig. 3A). This corroborates the metagenome results by Roco *et al*. [60] from acidic soils showing dominance of *qnor* over all other denitrification genes, suggesting that low pH selects for this gene. Interestingly, *cnor* apparently played a minor role in controlling NO, especially in Soil3.8 where gene abundance was close to 0 and transcripts were almost undetectable (Fig. 3). The taxonomic analysis of *qnor* genes and transcripts (Fig. 6) demonstrated that this gene, similarly as *nirK,* is widely distributed over different phyla, with the largest representation in the *Proteobacteria, Actinobacteria, Bacteroidetes, Chlamydiae, Acidobacteria* and *Planctomycetes*. qNor is the product of a single structural gene, *norB*, existing alone or in a small operon [81], and it is conceivable that this gene/gene cluster is more readily transferred horizontally than the bigger *cnor* operon. Furthermore, recent evidence suggests that qNor is electrogenic [82], as opposed to cNor, and it can therefore be speculated that it provides an extra, energetic advantage to its host organism. Taken together, it is not unreasonable to conclude that, between the two functionally redundant NO reductases related to denitrification, it is qNor that plays the major role in controlling NO concentrations under denitrifying conditions, at least in these soils. A primer-based study would most likely not have detected this since current primers for *nor* mainly target proteobacteria, which severely biases the results, especially for *qnor* [42].

Nitrous acid (HNO_2_) in its gaseous form HONO plays a key role in atmospheric reactions by being a precursor for hydroxyl (OH) radicals, which in turn take part in the formation of undesired O_3_ in the troposphere. Emissions of HONO and their connection to biological N-transformations have gained increasing interest during the past years. It has only recently been shown that biological processes producing NO_2_^−^ in soils contribute substantially to HONO emissions [6, 83, 84]. However, while Su *et al.* [83] stated that the low NO_2_^−^ concentrations in acidic soils are consistent with loss of HNO_2_ as HONO, Oswald *et al*. [6] instead showed that soils with neutral and slightly alkaline pH generally emitted more HONO than acidic soils. Our results show more than 10 times higher HONO production from Soil3.8 compared to Soil6.8 (Fig. 2), but we also show that the major portion of the NO_2_^−^ is not reduced and released as HONO, but instead reduced to NO and subsequently to N_2_O through denitrification.

The lower abundance of NOS genes and transcripts compared to NIR and NOR could, theoretically, be the cause of the low N_2_O reduction in Soil3.8, but it is unlikely to explain the nearly complete lack of N_2_O reduction in Soil3.8 during the first 35 h of incubation (Fig. 1A). This severely delayed N_2_O reduction in acidic soil corroborates other studies of soils from the same site and is in line with the growing evidence for a strong negative correlation between soil pH and N_2_O/(N_2_O+N_2_) product ratios[12, 33], suggested to be due to impaired maturation of the NosZ enzyme under acidic conditions (pH<6.1) [37–39]. The present study detected transcripts from both *nosZ* clades in Soil3.8, which suggests that the problem of producing functional NosZ under low pH conditions applies to both clades. Moreover, the taxonomic analysis showed that this problem is general to a diverse range of bacteria (Fig. 6), which adds new knowledge to earlier qPCR-based investigations in which taxonomy was not addressed [38, 39]. Although metatranscriptomes in Soil3.8 were only analyzed from the first 3 h of the incubation, the gas kinetics suggest that functional NosZ was not produced until after 25 h (Fig. 1B). In a natural situation the N_2_O produced in this time period would be emitted to the atmosphere. It can only be speculated why functional NosZ started to be produced after prolonged incubation. One reason could be that successful maturation took place somewhat stochastically, slowly building up a functional pool of NosZ. Another possibility is that populations of organisms such as the *Rhodanobacter* strains described by Lycus *et al.* [18] increased. These bacteria were shown to reduce N_2_O under acidic conditions, but not at neutral pH [18, 85].

Another important insight provided in the present study is the analysis of *nos* accessory genes and transcripts (Fig. 4) encoding enzymes involved in maturation of, and electron transfer to, NosZ. To our knowledge, such analyses have hitherto not been reported from soils. It is important to note, though, that since little is known about the protein function and phylogenetic characterization of these proteins the datasets used in this analysis are likely incomplete. Genes encoding NosR, which is involved in electron transfer to NosZ clade I [46] and NosL, a Cu chaperone that delivers Cu^+^ to the NosZ apo-protein of both clades [45], are of particular interest since the presence and transcription of these genes may hold a clue to understanding the severely delayed NosZ function in Soil3.8. The abundance of *nosR* gene reads (RPM values) in the MG of Soil3.8 was almost twice as high in Soil6.8, which is in agreement with the higher abundance of *nosZ* clade I in Soil3.8. The abundance of *nosL* was instead similar in the two soils, reflecting that this gene is part of the *nos* operon both in clade I and clade II organisms. The abundance of gene reads encoding the *nosDYF* cluster were also similar in the two soils. The function of the enzymes encoded by this gene cluster, which is part of the *nos* operon both in clade I and II organisms, is not completely elucidated. The *Y* and *F* genes are thought to encode an ATP-binding cassette (ABC) transporter, while NosD has also been suggested to be involved maturation of the CuZ site [45, 86, 87]. Taken together, gene reads for all the accessory *nos* genes analyzed in this study were detected in both soils, which was not unexpected.

Transcript reads encoding all accessory *nos* genes of clade I were detected in Soil3.8, which could be taken to indicate that the organisms in this soil had the tools in place for NosZ function, including *nosZ* transcriptional activation and electron transfer to NosZ (by NosR) and CuZ site maturation (by NosL). To conclude, there were no obvious issues with the genetic potential or the transcriptional activity of the *nos* operon which could explain the delayed N_2_O reduction in Soil3.8.

## Conclusions

Despite the vast number of denitrification studies, -omics based analyses are hitherto scarce. The present study added several pieces of information to the current understanding of soil denitrifier communities and how pH affects their activity, which organisms are involved, and their control and accumulation of denitrification intermediates. The metatranscriptomic results suggest that NO_3_^−^ reduction was dominated by NarG activity, with lesser contributions from NapA. The NO_3_^−^ reduction was apparently connected to denitrification and not to DNRA activity since N gas+ nitrosation accounted for ~100% of the consumption of NO_3_^−^. This is surprising and indicates a minor role of DNRA in N-transformations in these soils, despite DNRA-related gene and transcript reads (*nrfA* and *nirB*) being abundant.

The concentration of NO_2_^−^ in soil moisture was about two orders of magnitude higher in Soil6.8 than in Soil3.8, while the NAR/NIR transcript read ratios were almost identical in the two soils. This implies that NO_2_^−^ concentrations in Soil3.8 were controlled by a combination of biological activity and chemical degradation, which supports the experimental evidence from nitrite kinetics and modelling by Lim *et al*. [28]. It is conceivable that denitrifying organisms in acidic environments tune their genetic regulation to avoid high levels of NO_2_^−^, which at low pH will occur mainly as toxic HNO_2_, and at the same time avoid high NO concentrations by maintaining high transcription of NOR, mainly *qnor*, which was apparently selected for over *cnor* by low pH. Our finding that HONO emissions from Soil3.8 were 10 times higher than from Soil6.8 is noteworthy since, while it supports the results by Su *et al.* [83] it contradicts other published results which indicate that neutral HONO emissions are mainly from neutral soils [6]. More experimental evidence is however required to reach a better understanding of environmental conditions affecting the release of HONO from soils.

The analyses of the MG and MT demonstrated that diverse microorganisms transcribed *nosZ* genes from both clade I and II in the acidic soil, but this was not followed by any detectable Nos function until >30 h after the start of the incubation. The results did not reveal any obvious lack of genes or transcripts of *nos* accessory genes involved in NosZ maturation. Although not exhaustive, this information provides a new building block towards an understanding of the impaired N_2_O reduction under acidic conditions.

The discrepancies between qPCR and -omics based estimates of genes and transcripts provides strong evidence that primers commonly used in denitrification studies only capture a fraction of the community that carries these genes. The problem likely arises since genes such as *nirK* occur in diverse of microbes, while genes such as *nirS* are restricted to a narrower group of denitrifiers making the *nirS* primer sets better able to bind to the targeted gene sequences. This leads to an overestimation of *nirS* relative to *nirK* genes and transcripts in the qPCR-results. Similarly, the qPCR results showed lower abundance of *nos* (clade I) in Soil3.8 than in Soil6.8, while it was 3 times more abundant in the metagenome of Soil3.8 compared to Soil6.8. These examples illustrate that primer-based results for denitrification genes may lead to erroneous conclusions and must be interpreted with caution.

## Supporting information

Supplementary files_Frostegard et al

## Acknowledgements

SV was financed by the project PASUSI within the ERA-NET Co-fund HorizoN2020 program Leap-Agri, under the EC Grant agreement No. 727715 and the Research Council of Norway projects No. 290488 and No. 286888. NL obtained a PhD grant from the Norwegian University of Life Sciences.

