## Supplementary files_Frostegard et al for "Linking meta-omics to the kinetics of denitrification intermediates reveals pH-dependent causes of N_2_O emissions and nitrite accumulation in soil"

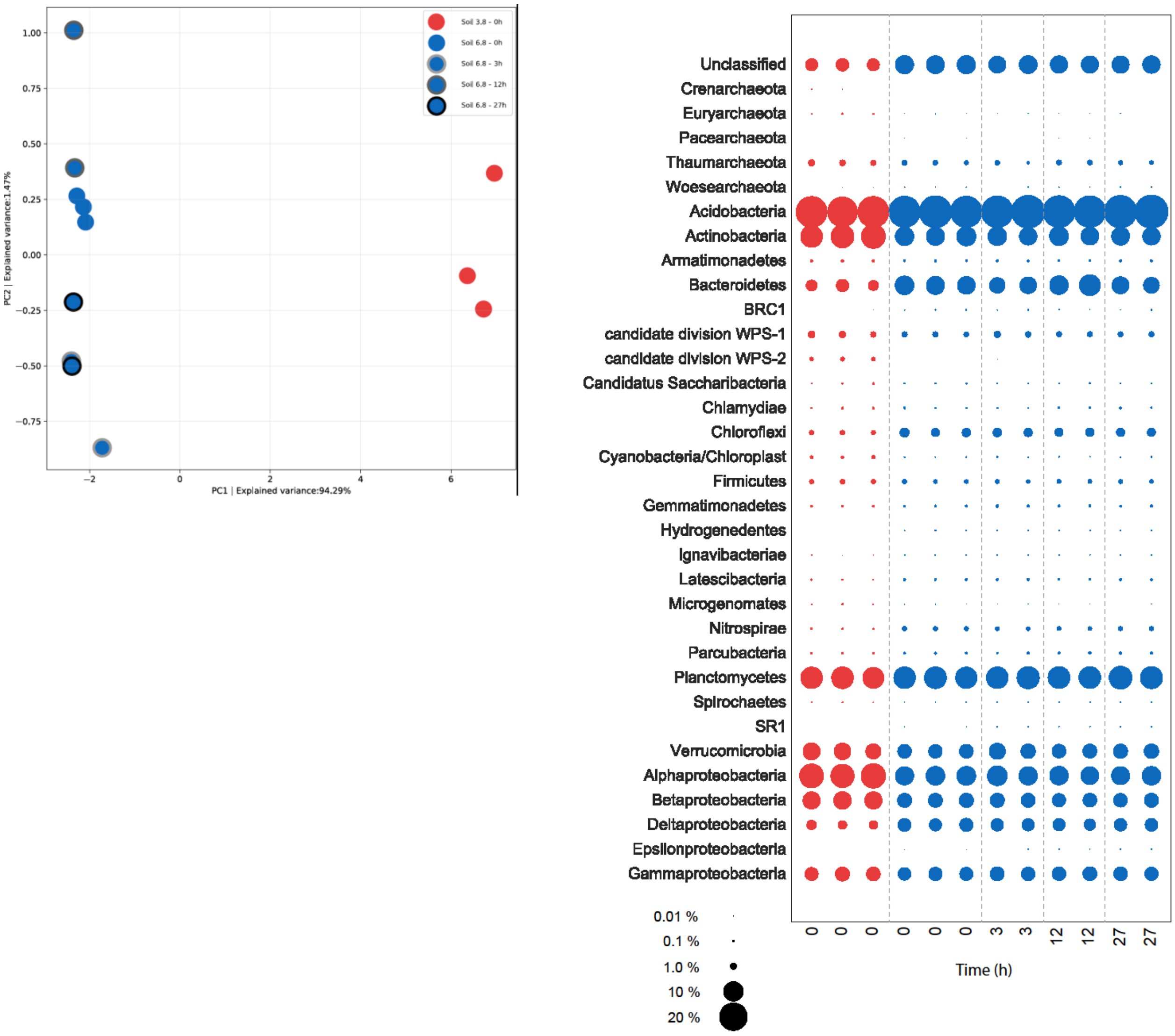


**Fig. S1.** Taxonomic distribution of bacteria in SoilpH 3.8 and SoilpH 6.8 based on 16S rRNA gene sequences. A) Principal component analysis (PCA). B) Taxonomic breakdown. Initial time point samples (0 h incubation, both soils; n=3). For Soil6.8 these were compared against grouped later time point samples taken after 3, 12 and 27 hours of incubation (n=2). Samples were sequenced on an Illumina MiSeq platform after amplification of the V4 region of the 16S rRNA gene (primers 515f/806R).

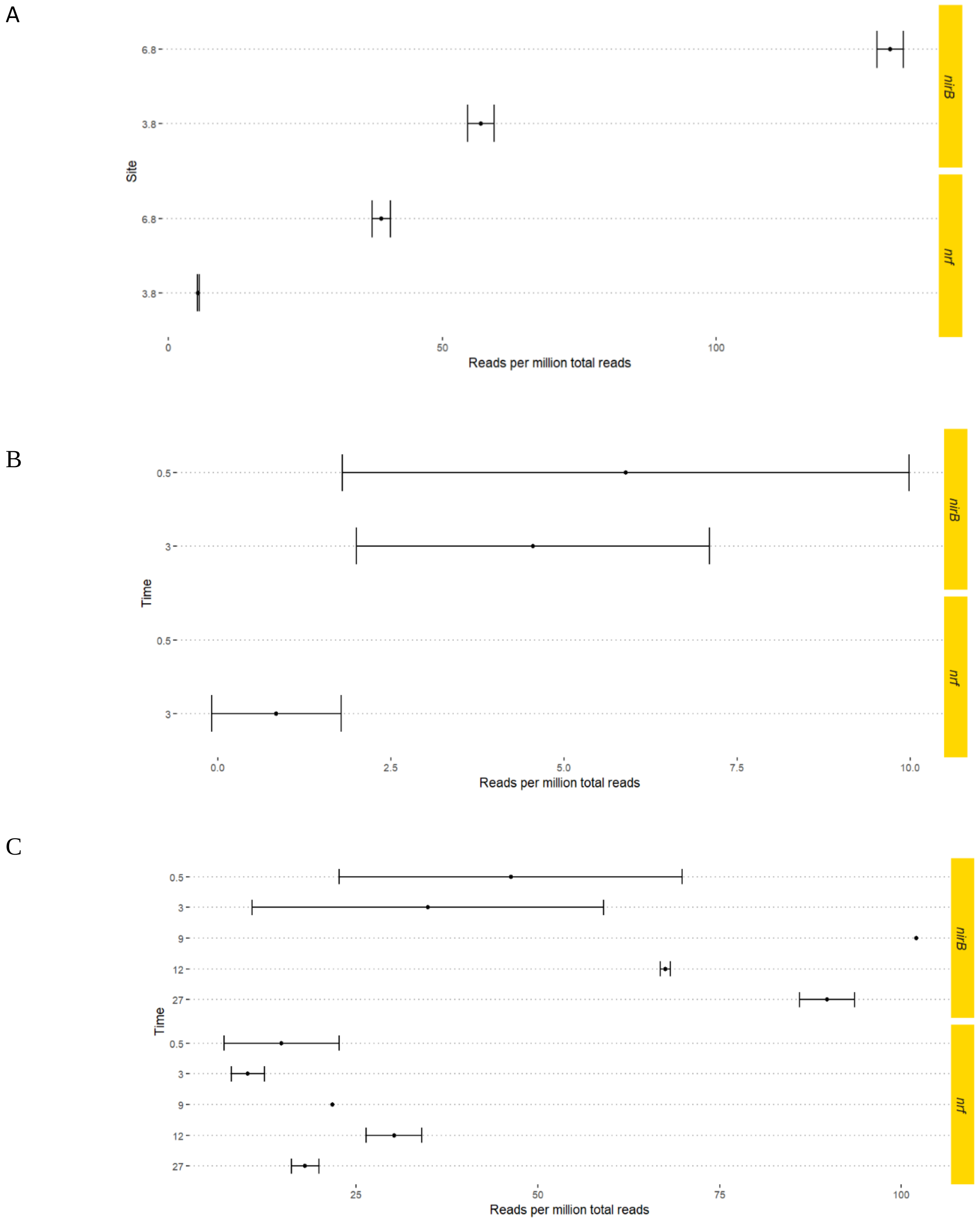


**Fig. S2.** Genetic potential and transcription of the DNRA-related genes *nrfA* and *nirB* in the metagenome and metatranscriptome of Soil3.8 and Soil 6.8. A) Average gene read abundances (as reads per million, RPM). Bars show sd, (n=3). B) Average transcript read abundances after 0.5 and 3h (Soil3.8). C) Average transcript read abundances after 0.5, 3, 9, 12 and 27 h of incubation (Soil6.8). B and C (transcript reads): Duplicate samples were analyzed for each sampling point, bars show highest and lowest value.

**Calculating the concentration of $HNO_2$ as a function of pH and 31 total nitrite concentration.**

The Henderson-Hasselbalch equation (1) can be used to calculate the relative amount of undissociated nitrite as a function of pH (which is controlled by the buffer system of the soil):

$K_a = [H^+]*[A^-]/[HA]$ (1)

Where $K_a$ is the base dissociation constant

Taking the $\log_{10}$ of both sides

$\log_{10}(K_a) = \log_{10}[H^+] + \log_{10}([A^-]/[HA])$ (2)

defining pX = - $\log_{10}$[X], (2) gives:

$-pK_a = -pH + \log_{10}([A^-]/[HA])$ (3)

Replacing $[A^-]$ with $[NO_2^-]$ and [HA] with $[HNO_2]$,
and solving equation (3) for $[NO_2^-]/[HNO_2]$:

$[NO_2^-]/[HNO_2] = 10^{(pH-pKa)}$ (4)

Equation (4) can be solved for $[HNO_2]/([HNO_2]+[NO_2^-])$, which is the molar fraction of total nitrite (as measured) that is un-dissociated:

$[HNO_2]/([HNO_2]+[NO_2^-]) = 1/(1+[NO_2^-]) = 1/(1+10^{(pH - pKa)})$ (5)

Hence, we can calculate the concentration of un-dissociated $HNO_2$ in the soil
$[HNO_2] = TNN/(1+10^{(pH - pKa)})$ (6)

where TNN is the measured concentration of total nitrite N ($[HNO_2]+[NO_2^-]$, pH is the measured soil pH and $K_a$ is the dissociation constant for nitrous acid, which is 4E-4 (hence $pK_a$=3.3398).

Needless to say, soil pH is the most problematic parameter, since the pH may vary throughout the soil matrix, and the bulk pH as measured depends on the cation concentration in the soil slurry. Our pH measurements were done in 0.01 M $CaCl_2$, which is thought to give pH values close to the average of the intact soil. Higher salt concentrations will give lower pH values and *vice versa*.
